# Promoter architecture determines co-translational regulation of mRNA

**DOI:** 10.1101/192195

**Authors:** Lorena Espinar, Miquel Àngel Schikora Tamarit, Júlia Domingo, Lucas B. Carey

**Author notes:** Corresponding author. (L.B.C.). Equal contribution (Lorena Espinar, Miquel Àngel Schikora Tamarit, Júlia Domingo).

## Abstract

Information that regulates gene expression is encoded throughout each gene but if different regulatory regions can be understood in isolation, or if they interact, is unknown. Here we measure mRNA levels for 10,000 open reading frames (ORFs) transcribed from either an inducible or constitutive promoter. We find that the strength of co-translational regulation on mRNA levels is determined by promoter architecture. Using a novel computational-genetic screen of 6402 RNA-seq experiments we identify the RNA helicase Dbp2 as the mechanism by which co-translational regulation is reduced specifically for inducible promoters. Finally, we find that for constitutive genes, but not inducible genes, most of the information encoding regulation of mRNA levels in response to changes in growth rate is encoded in the ORF and not in the promoter. Thus the ORF sequence is a major regulator of gene expression, and a non-linear interaction between promoters and ORFs determines mRNA levels.

## Introduction

Precise control of gene expression is essential (Skotheim et al. 2008) and the way this regulatory information is encoded in enhancers, promoters, 5’ untranslated regions (UTRs) and other regulatory regions throughout the gene sequence has been well characterized (Shalem et al. 2013; Keren et al. 2013; Dvir et al. 2013; Kudla et al. 2009; Shalem et al. 2015). Steady-state mRNA levels are determined by the combination of synthesis and decay rates. It was generally assumed that promoters mostly determine synthesis rates, while sequences in the 5’UTR, open reading frame (ORF) and 3’UTR determine mRNA translation and degradation rates. While there is now evidence that synthesis, decay and translation are tightly coupled,there is no clear delineation of regulatory mechanisms within different parts of a gene and the precise mechanisms that link the promoter to cytoplasmic processes remain unknown (Haimovich et al. 2013; Wolffe and Meric 1996; Ladomery 1997).

As the location that regulates the initiation of transcription, the promoter is a reasonable place to start thinking about mRNA expression. In yeast, genes can broadly be split into two classes based on promoter architecture, those with a consensus and evolutionarily conserved TATA box, and those without one (TATA-less). TATA-binding protein (TBP) binds TATA+ promoters at the TATA box, but TBP also binds TATA-less promoters at a TATA-like sequence that is one or two mismatches away from the TATA consensus. TATA-less promoters are TATA-like. While recent work suggests that this distinction is not quite binary, multiple lines of evidence have shown that, on many different levels, genes which lack a consensus evolutionarily conserved TATA box are fundamentally different from the TATA+ class (Taatjes 2017; Kubik et al. 2017). Genes lacking a TATA box are depleted of nucleosomes around the transcription start site (TSS) and exhibit lower transcriptional plasticity (Tirosh and Barkai 2008; Rhee and Pugh 2012). For simplicity, we will generally refer to the TATA+ class as constitutive and the TATA-like (TATA-less) class as inducible.

Features within the open reading frame itself, such as codon usage, correlate with steady-state mRNA levels (Akashi 2003; Neymotin et al. 2016, 2014). There are two proposed reasons for this correlation: natural selection for specific codon usage and an active role in codon usage in regulating mRNA levels. In first, codon usage is shaped by ribosome dynamics and tRNA pools, with selection for accurate and fast translation, especially among abundant proteins (Plotkin and Kudla 2011). In the second, codon usage directly affects mRNA levels (Chen et al. 2017), likely through translation-coupled decay. There is ample evidence for both models, suggesting a model in which both the act of translation and selection on codon usage are together responsible for the observed correlation between codon usage and gene expression among native genes.

In addition to effects from codon usage, the stability of transcripts is determined by multiple other processes, one of which is nonsense mediated decay (NMD). Initially identified as a quality control mechanism for transcripts that contain an aberrant premature termination codon (PTC) within the ORF, there is now abundant evidence that NMD affects the expression of between 5% and 20% of native transcripts in yeast and mammalian cells (Behm-Ansmant et al. 2007; Kervestin and Jacobson 2012; He and Jacobson 2015; Tani et al. 2012). The precise mechanism is still unclear, but three models have been proposed: the Exon Junction Complex (EJC) model, the Upf1 3’-UTR sensing and potentiation model, and the faux 3’-UTR model (He and Jacobson 2015; Le Hir et al. 2000; Amrani et al. 2004; Hogg and Goff 2010). Previous observations suggest that the EJC model is the main source for NMD in mammals, but yeast mainly trigger NMD through the other two mechanisms (Lindeboom et al. 2016). The latter have a common denominator: long 3’UTRs, which result from PTCs but can also be found in native mRNAs. Thus, 3’UTR length encodes regulatory information in a way that depends on active translation.

To further complicate the relationship between sequence and expression, some transcription factors (TFs) influence mRNA stability and localization (Braun et al. 2015; Bregman et al. 2011; Zid and O’Shea 2014). Thus, in addition to affecting mRNA synthesis rates, promoters play a role in other parts in the life of an mRNA. However, three major questions remain: by which mechanisms do elements in the promoter influence the lifecycle of the mRNA, where in the gene are different types of regulatory information encoded, and are there genetic interactions between discrete regulatory regions?

Promoters influence mRNA levels. Coding sequences influence mRNA levels. To determine if there is a genetic interaction between these two spatially distinct regulatory regions, we generated a pooled library of over 10,000 yeast genomic DNA fragments cloned into plasmids containing either inducible TATA box containing GALL promoter or the constitutive TATA-less RPL4A promoter. This library exhibits over four orders of magnitude of expression, most of which can be predicted using a sequence-feature based mathematical model. Intriguingly, in both the library and in native transcripts, co-translational features such as codon usage more strongly influence mRNA levels when expression is driven by a constitutive promoter. To identify the molecular mechanism for this difference in co-translational regulation between promoter architectures we performed a computational genetic screen across 6402 RNA-seq experiments, and found that the RNA helicase Dbp2 specifically insulating transcripts of TATA+, but not TATA-less promoters from the effects of co-translational regulation on steady-state mRNA levels. Finally, we used RNA-seq and promoter-YFP expression data to show that, specifically for TATA-less promoters, the ORF contains more regulatory information than the promoter.

## Results

### ORF-encoded sequence features play an active role in regulating gene expression in native transcripts

To determine the ability of coding sequences to regulate gene expression we developed a method to decouple native genomic sequences from their 5’ and 3’ regulatory context (**Fig. 1A, Supplemental Fig. 1**). We digested the yeast genome with four restriction enzymes and inserted the resulting random genomic DNA fragments into a plasmid containing the inducible GALL promoter, a start codon, the ClaI restriction site and three staggered stop codons (**Supplemental Fig. 1A**). This generated a pool of over 10,000 plasmids, each of which contains a single random fragment of the yeast genome. The cloned random gDNA fragments derive from the entire yeast genome and therefore contain parts of native coding sequences as well as untranslated regions (UTRs), promoters, transcription terminators and other chromosomal features (**Supplemental Fig. 1B, Supplemental Table 1**). We transformed the plasmids into yeast and measured expression of each transcript as the ratio between RNA and DNA abundance (**Fig. 1B**). In spite of having the same promoter, 5’ untranslated region and transcription terminator, the expression of the gDNA fragments varies of over four orders of magnitude (**Fig. 1C, Supplemental Fig. 2**). To determine the sources of this regulation and to quantify the contribution of individual sequence features to changes in expression, we computed over 4000 sequence features for each insert, such as dimer and trimer nucleotide counts, GC content, codon bias and the presence of premature stop codons (**see methods**). We fit a linear model with a minimal number of predictive features to the experimental data and found that a model with seven features can accurately predict expression (R^2^=0.64 for all inserts, and R^2^=0.61 for inserts with no premature stop codon) (**Fig. 1D, Fig. 2B, Supplemental Fig. 3**), suggesting that a small number of ORF-encoded sequence features generate a large amount of the variation in gene expression. To determine if ORF-encoded sequence features regulate expression in native genes we used the same model, without taking into account 3’UTR length to predict expression of native genes from their coding sequence (**Fig. 1E**).

**Figure 1.**
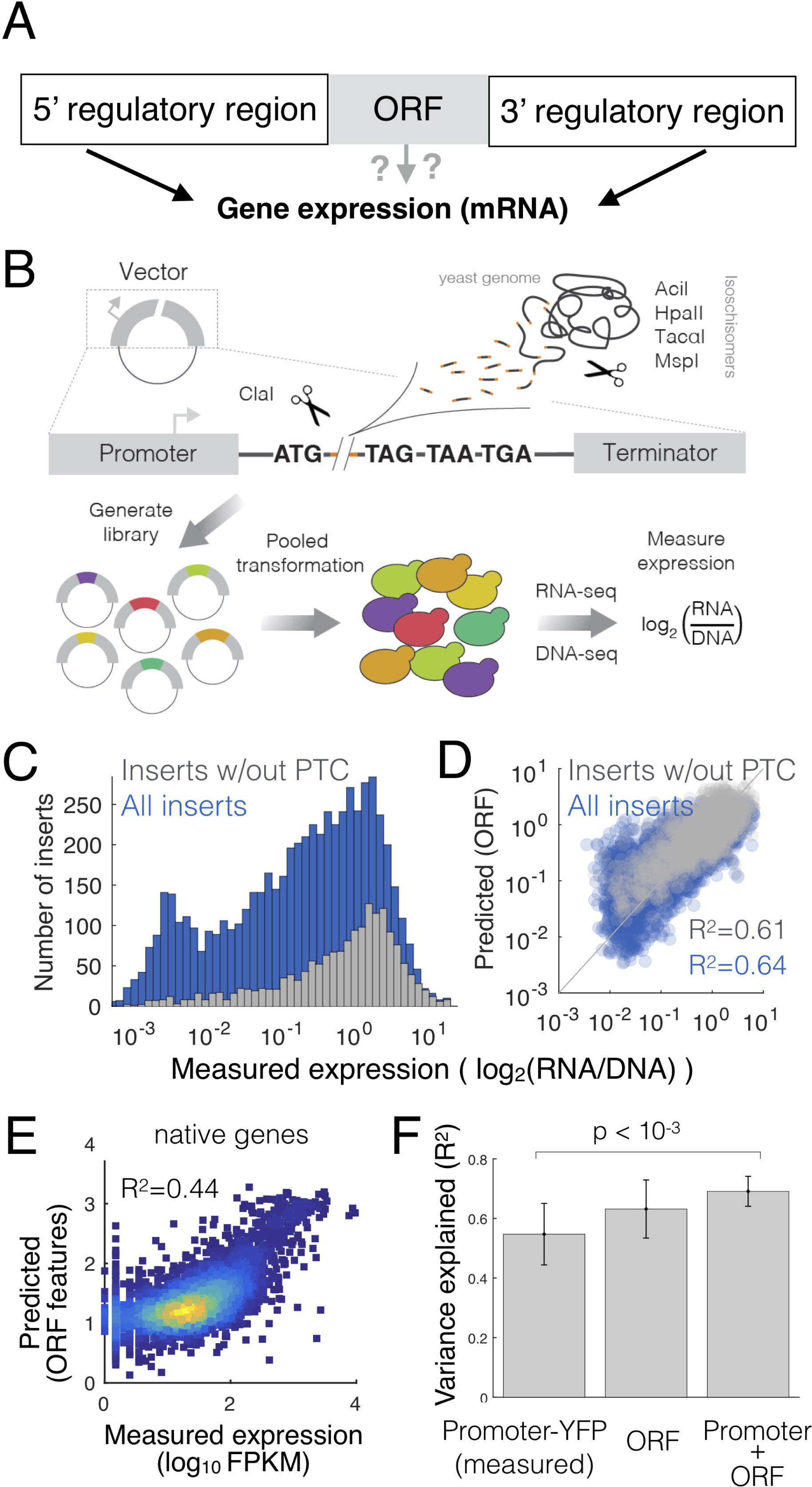
An expression library of genomic fragments to quantify the ability of ORF-encoded sequence features to regulate gene expression. (**A**) A schematic view of the initial question: *Does the open reading frame determine gene expression?* (**B**) Scheme of the gDNA library preparation and expression measurements. (**C**) Measured expression distributions for all inserts (blue) and only those lacking a premature termination codon (grey). (**D**) Measured vs predicted expression levels in the gDNA library. Expression is predicted from the sequence of each gDNA insert using a 10-fold cross-validated linear model (R^2^ is calculated across all test data from all cross-validations). (**E**) Expression predicted from the sequence of each native yeast ORF using the same features as for the gDNA library. (**F**), Including ORF-encoded features in a model of expression increases the ability of promoter-YFP data to predict steady-state mRNA levels. Error bars are standard deviation from 10-fold cross-validation.

**Figure 2.**
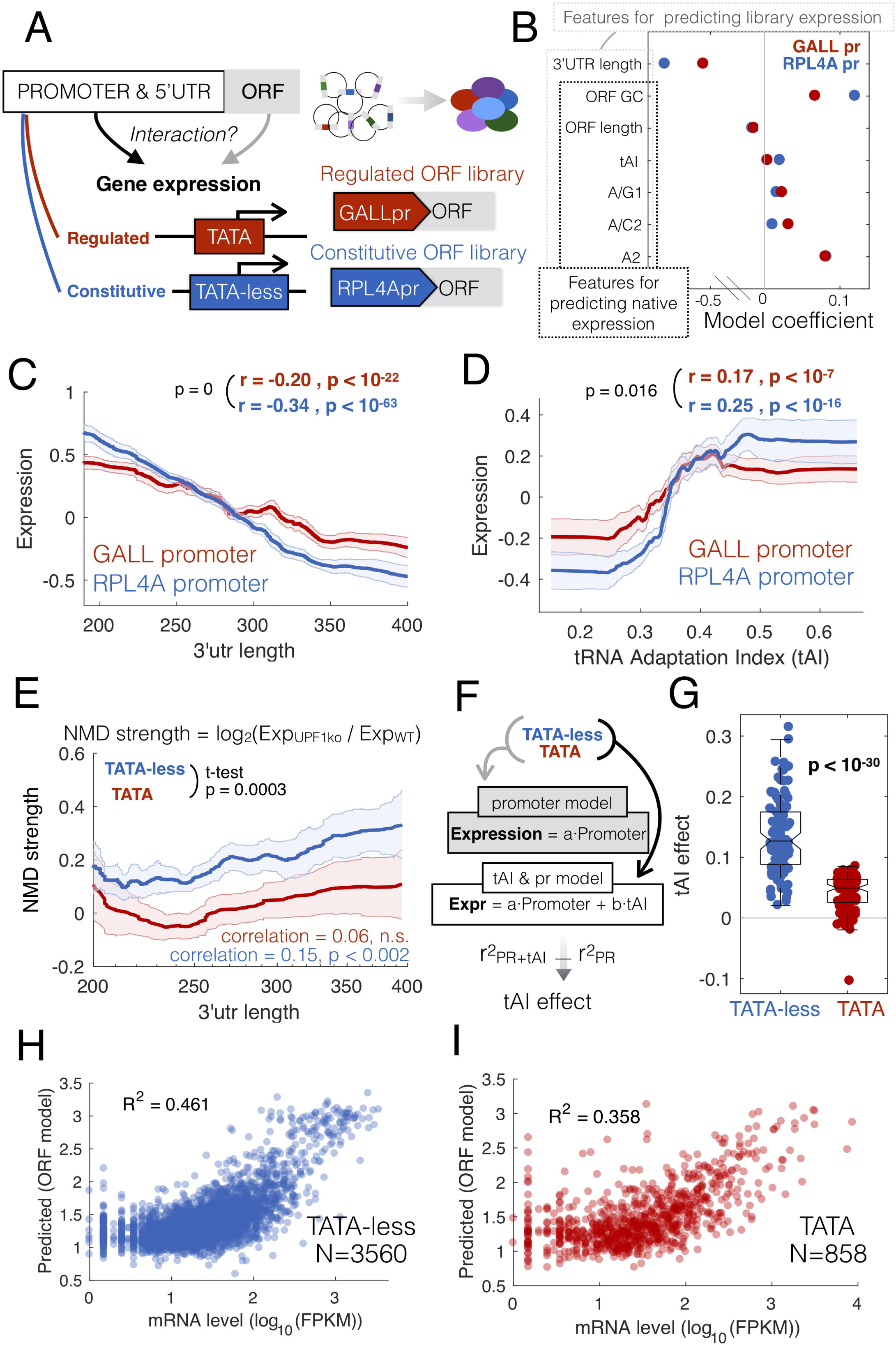
A gDNA library with different promoters identifies sequence features that interact with the promoter to determine gene expression. (**A**) The gDNA library was cloned under the control of either the TATA+ GALL promoter or the TATA-less ribosomal RPL4A promoter and the expression of both libraries measured in yeast growing on galactose as a carbon source. (**B**) Coefficients from the multiple linear model based on ORF sequence features from libraries with the two different promoters. Outlined are the features used for predicting expression in both the libraries or native genes. tAI is the tRNA adaptation index. Nucleotides followed by numbers refer to the position in a codon, eg: A/G1 is the fraction of codons with an A or G at position 1. (**C,D**) Lines show the median expression for inserts binned by 3’UTR length (c) or codon bias (d). Correlation values are for unbinned data and the p-value is a test for a significant difference between the two correlation values using bootstrapping. (**E**) NMD effect, measured as the log2 ratio in mRNA (TPM) between *upf1* and wild-type cells for native transcripts. Lines show the median NMD effect across transcripts binned by 3’UTR length for TATA-containing (red) and TATA-less promoters (blue). The p-value is for a t-test for a difference in mean NMD strength for all unbinned data between TATA and TATA- less genes. (**F**) The makeup of two linear models, one that predicts mRNA levels from promoter-YFP data, and the other that includes codon bias (tAI) as an additional predictor. For both models, native genes are split into two classes, TATA and TATA-less, and tAI effect is the difference in R^2^ between the two models. (**G**) Difference in tAI effect for random samplings of equal numbers of genes from each class. (**H,I**) An ORF-encoded sequence feature model was trained to predict mRNA levels for TATA and TATA-less promoters (R^2^ = squared Pearson correlation coefficient).

The ability of the ORF-feature model to predict expression of native genes could be because coding sequences play an active regulatory role, or it could be due to co-evolution and selection for certain sequence properties in genes whose expression levels are determined by more stereotypical regulatory regions such as the promoter. To differentiate correlation from causation we took advantage of data in which the strength of 859 promoters driving YFP was measured (Keren et al. 2013). We find that ORF features are equally good as promoters in predicting steady-state mRNA levels, and that a combined model including both promoter-YFP data and ORF features performs significantly better (**Fig. 1F**). Thus, ORF-encoded sequence features play an active role in regulating gene expression in native transcripts.

### Co-translational regulation is stronger for mRNAs regulated by TATA-less promoters

While the majority of experiments that investigated transcriptional, co-translational and post-translational regulation used inducible promoters (Shalem et al. 2013; Puchta et al. 2016; Meaux et al. 2008; Shah et al. 2013; Radhakrishnan et al. 2016), most of these promoters are in fact not condition-specific. Promoters can be roughly divided into two broad categories, those with a TATA box (SAGA-dominated and inducible or regulated) and those lacking a conserved TATA box (TFIID-dominated and constitutive) (Basehoar et al. 2004; Tirosh and Barkai 2008; de Jonge et al. 2016; Struhl 1986; Huisinga and Pugh 2004). To determine if ORF-encoded sequence features are regulatory for constitutive promoters we built the random gDNA fragment library in a second plasmid in which the regulated TATA box containing GALL promoter was replaced with the equally strong constitutive ribosomal RPL4A promoter (**Fig 2A, Supplemental Fig. 4**). We found that codon bias (tRNA Adaptation Index (tAI) (dos Reis et al. 2003)) and 3’UTR length, both of which require active translation to be interpreted by the cell (Gardin et al. 2014; Kervestin and Jacobson 2012; Presnyak et al. 2015), have more importance in a model trained on RPL4Apr data compared to GALLpr data (**Fig. 2B**). Both 3’UTR length and codon bias have a stronger effect on expression when transcription is driven by the constitutive RPL4A promoter (**Fig. 2C,D, Supplemental Fig. 5**).

To determine if this promoter-specific effect of co-translational regulation affects native genes we divided promoters into two categories, those with an evolutionarily conserved TATA box (inducible) and those without (TATA-less, or constitutive) (Basehoar et al. 2004). To measure the effect of NMD separately for each class of transcript we used RNA-seq measurements from a Δ*upf1* strain, which is defective in nonsense-mediated-decay (NMD) (Smith et al. 2014). Consistent with the gDNA library, native transcripts from TATA-less promoters are affected by NMD in a manner that increases with 3’UTR length, while transcripts from promoters with an evolutionarily conserved TATA box are unaffected by NMD (**Fig. 2E, Supplemental Fig. 6**). To determine if the effect of codon bias on native genes is stronger for TATA-less promoters we used a linear model to predict mRNA levels from either promoter-YFP expression data alone (Keren et al. 2013) or from a model that includes both promoter-YFP data and codon bias (tAI (dos Reis et al. 2003)) and measured the increase in R^2^ when including codon bias as a model feature (**Fig. 2F**). Consistent with results from the gDNA library, codon bias is more strongly predictive of mRNA levels for transcripts driven by TATA-less promoters (**Fig. 2G**). These differences are not due to differences in expression, mRNA stability or codon bias, since the distribution of mRNA expression, degradation rates and tAI are similar between the two classes of genes (**Supplemental Fig. 4**). To determine if ORF-encoded sequence features are more predictive of native expression levels for TATA-less genes, we predicted expression from sequence separately for the two classes of genes. ORF features are more predictive of expression in TATA-less genes (**Fig. 2H,I**). In both, for the gDNA library and native genes, co-translational regulation affects mRNA levels more strongly when expression is driven by TATA-less promoters, suggesting the existence of some mechanism that carries information from the promoter to the ribosome.

### A genetic screen of 6402 RNA-seq experiments identifies the RNA helicase Dbp2 as TATA containing promoter specific insulator damper of co-translational regulation

To identify the molecular mechanism underlying the difference in co-translational regulation between TATA and TATA-less promoters we performed a computational genetic screen for mutants that alter our ability to predict mRNA levels from promoter-YFP data. We screened 6402 RNA-seq experiments, all RNA-seq experiments that have been done in *S. cerevisiae (Ziemann et al. 2015)*, for mutants that affect the ability of codon bias to predict expression in a promoter-class-specific manner (**Fig. 3A**). For each of the 6402 experiments we calculated the increase in R^2^ upon including codon bias as a predictive feature. Codon bias has more predictive power for TATA-less genes across thousands of RNA-seq experiments (**Fig. 3B**). We identified a single mutant which significantly reduces this difference across multiple biological replicates. Deletion of Dbp2 (Beck et al. 2014) increases the effect of codon bias on gene expression specifically for TATA containing genes (**Fig. 3C, Supplemental Fig. 7**). Furthermore, DBP2 expression varies across the dataset and this variation correlates with the effect of tAI. Conditions and mutants with low DBP2 expression have a higher tAI effect on expression of TATA+ but not TATA-less genes (**Supplemental Fig. 7A**). This suggests that Dbp2, a co-transcriptionally loaded RNA helicase (Ma et al. 2016; Cloutier et al. 2012), specifically reduces the effect of co-translational regulation on mRNA levels in a promoter-specific manner. Interestingly, Dbp2 physically interacts with nucleosomes, Pol-II-associated GTFs, and the ribosome (Stark et al. 2006) (**Supplemental Fig. 8**).

**Figure 3.**
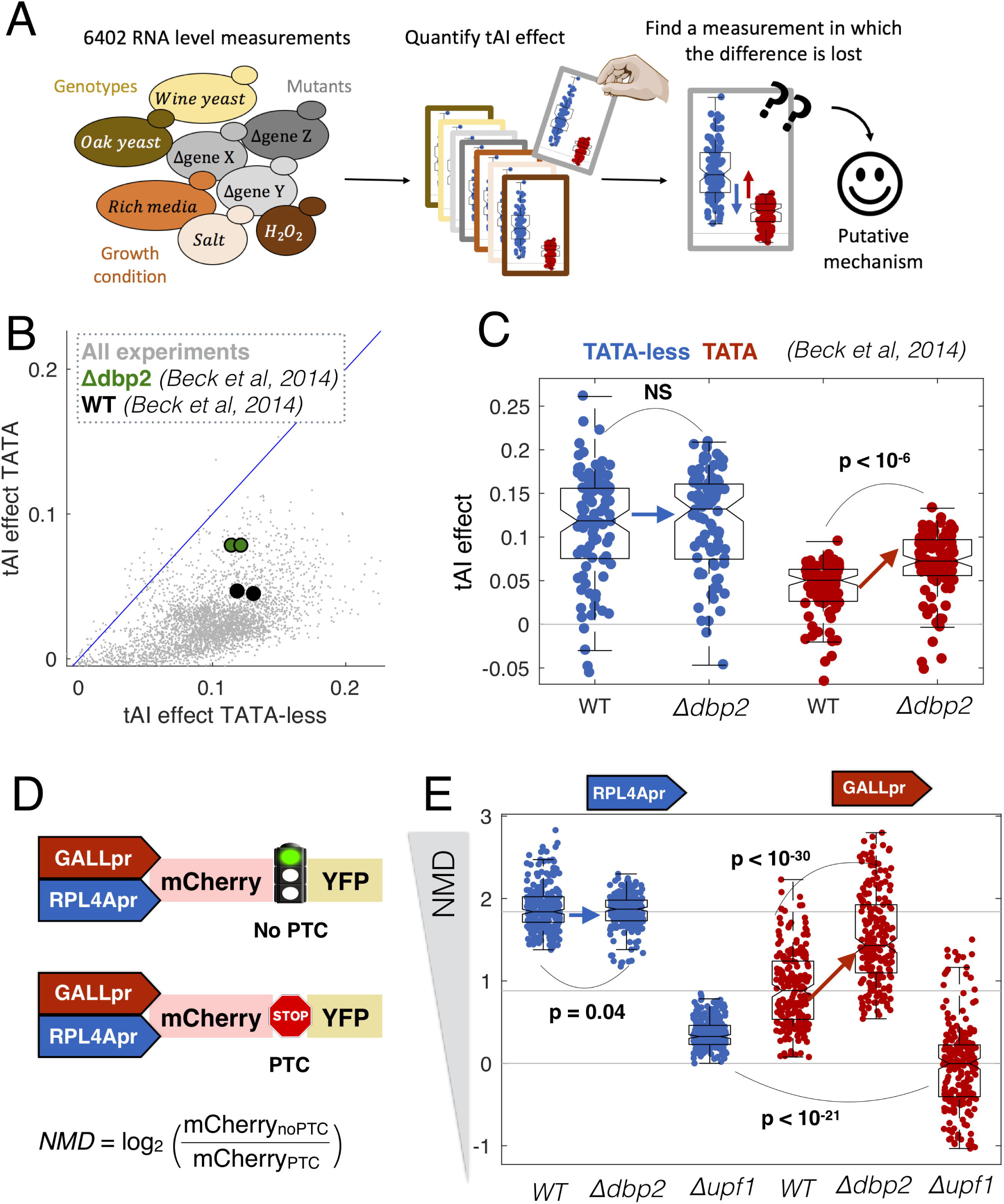
A computational genetic screen for mutants that alter promoter-specific co-translational regulation. (**A**) We analyzed expression data for 6402 RNA-seq experiments in *S. cerevisiae*, and for each experiment predicted mRNA levels from promoter-YFP data alone, or promoter-YFP and codon bias (tAI). (**B**) For each for the 6402 experiments we calculated the ability of tAI to improve R^2^ for TATA-less and TATA-containing genes. Highlighted is a single experiment (Beck et al. 2014) in which the R^2^ changes in a promoter-class-specific manner between wild-type and mutant cells. (**C**) Shown are the changes in R^2^ for wild-type and Δ*dbp2* cells for TATA containing and TATA-less transcripts. Each point corresponds to the effect of tAI on expression from equal-sized random samples of TATA (red) and TATA-less (blue) genes. Boxplots show the median and interquartile range. p-values are from t-tests. (**D**) Synthetic system for measuring the effect of NMD on gene expression. NMD is the ratio in mCherry expression between plasmids with and without a premature termination codon (PTC). (**E**) In each genotype we measured mCherry expression 16 times (biological replicates) for each of the four plasmids. As NMD effect is the ratio between two plasmids, for each promoter and genotype we calculated 8^2^ ratios. Boxplots show the median and interquartile range for each set of ratios.

To confirm the results of the screen and to determine if Dbp2 plays a role in other forms of co-translational regulation of mRNA levels, such as NMD, we built a set of four synthetic NMD reporters (**Fig. 3D**). Each plasmid contains two fluorescent proteins, mCherry and YFP, with or without a premature termination codon (PTC) in the intervening linker. In the presence of a stop codon in the linker, YFP becomes a long 3’UTR and this targets the transcript for NMD (Muhlrad and Parker 1999). Each of the two constructs is driven by either a constitutive (RPL4A) or a regulated (GALL) promoter. We transformed the plasmids into wild-type, *Δupf1* and *Δdbp2* yeast and measured mCherry and YFP expression (**Supplemental Fig. 4**). The NMD effect is the ratio of the mCherry signal in the noPTC and PTC constructs. In wild-type cells the effect of NMD is stronger in the TATA-less RPL4Apr plasmids, consistent with the gDNA library and native genes. Consistent with the computational RNA-seq screen, transcripts from the inducible GALL promoter are more strongly affected by NMD in a *Δdbp2* strain (**Fig. 3E**). This suggests a model in which Dbp2 mutes the effect of codon bias and NMD on mRNA levels specifically for TATA containing promoters.

### For TATA-less promoters, growth-rate mediated regulation of expression is implemented in the coding sequence, not the promoter

Why implement regulation in coding sequences? In response to changes in growth rate, cells alter the expression of thousands of genes (Brauer et al. 2008; García-Martínez et al. 2016). In yeast, expression of translation-related genes is positively correlated with growth rate, while genes associated with environmental stress response, respiration and oxidative phosphorylation are negatively correlated with growth rate (**Fig. 4A, Supplemental Table 2**)(Brauer et al. 2008). TATA-less genes tend to be positively correlated with growth rate, while TATA containing genes are negatively correlated (Fisher’s exact test p=10^−27^, odds ratio = 4.0). To determine if the information encoding regulation of expression in response to changes in growth rate is in the promoter, we took advantage of a dataset in which promoter-YFP expression data for 859 genes were measured in ten different environmental conditions (Keren et al. 2013). For each gene we calculated the slope, how promoter-YFP expression changes with growth rate (**Fig. 4A**), and compared it to the change in steady-state mRNA levels upon changes in growth. We find that changes in the mRNA levels of constitutive genes (TATA-less) are worse predicted by promoter-YFP data (**Fig. 4B**). Therefore, for TATA-less genes, information encoding growth-rate mediated regulation of expression lies outside of the promoter. To determine if the information that regulates expression with growth rate is encoded in the ORF, we asked if ORF encoded sequence features can predict how mRNA levels change as a function of growth rate. We find that ORF features predict growth-mediated changes poorly for TATA genes and better for TATA-less genes (**Fig. 4C**). For constitutive genes, there is more regulatory information in the ORF than in the promoter (**Fig. 4D,E**).

**Figure 4.**
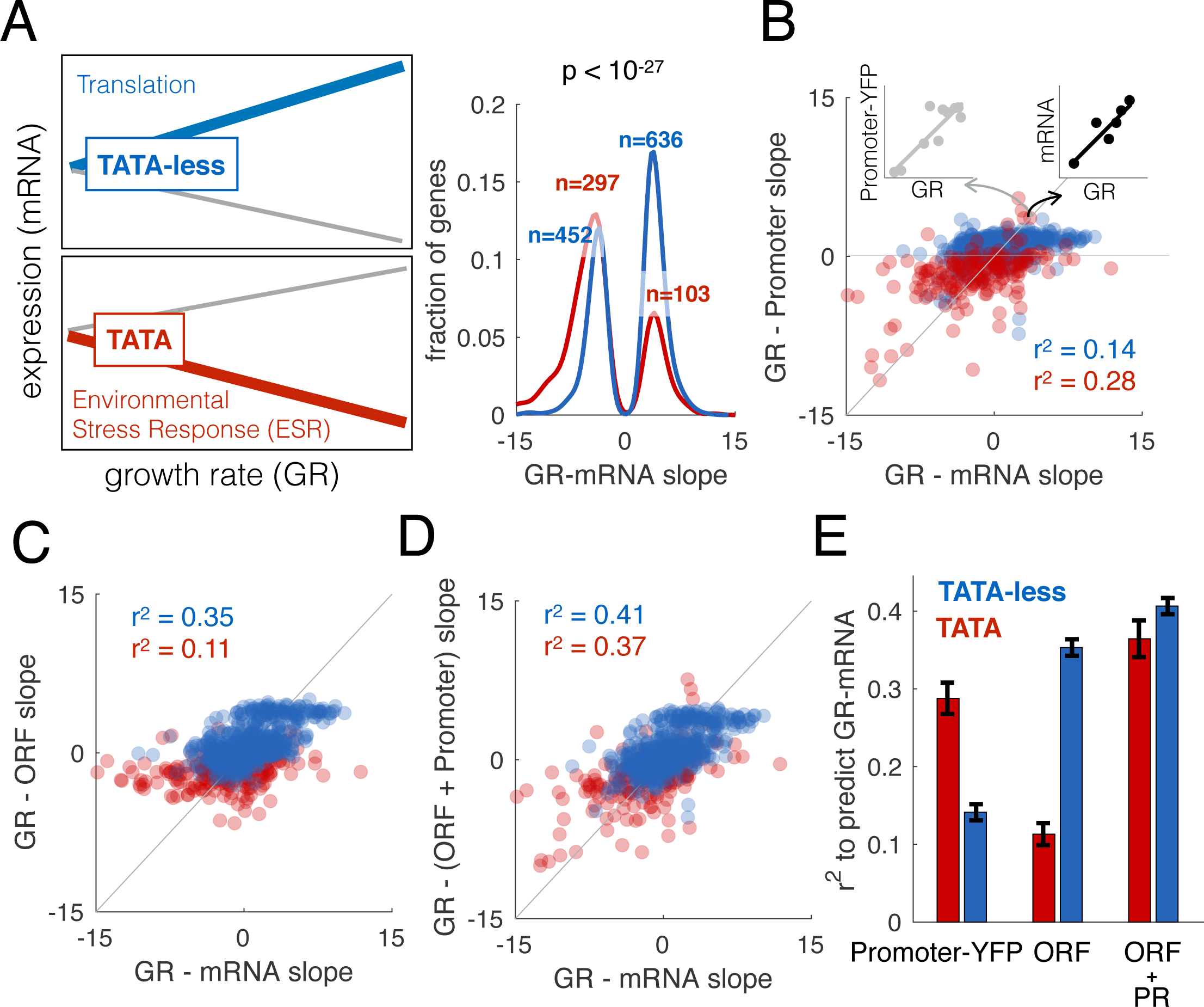
ORF features regulate expression in response to changes in growth rate. (**A**) Brauer et al(Brauer et al. 2008) grew yeast at six different growth rates in each of six different environments and measured how the expression of each gene changed as a function of growth rate (slope GR- mRNA). TATA-less genes (blue, N=1088) increase in expression, while those with TATA boxes (red, N=401) more often decrease. The p-value is from a Fisher’s Exact test. (**B**) Keren et al(Keren et al. 2013) measured promoter-YFP expression at different growth rates. Each point shows the slope (expression change as a function of growth rate) for promoter-YFP (y-axis) and steady-state mRNA (x-axis) for TATA (red) or TATA-less (blue) genes. (**C**) GR-mRNA was predicted from ORF features allowing us to calculate the slope between growth rate and the contribution of ORF to expression (GR-ORF). Shown is the relationship between GR-ORF and GR-mRNA (see fig. A), for each type of genes. (**D**) GR-mRNA was predicted from both GR-Promoter and ORF features (GR- (ORF + Promoter)). Shown is the relationship between this and GR-mRNA, for each type of genes (**E**) R^2^ values for a model that predicts GR-mRNA from GR-Promoter, ORF features or ORF + Promoter, for each type of genes. Error-bars correspond to the standard deviation, calculated with bootstrapping.

## Discussion

We used a pooled library of random mRNAs under the control of either the TATA+ inducible GALL promoter or the TATA-less constitutive RPL4A promoter to identify features inside of transcripts that exhibit quantitatively different effects on steady-state mRNA levels as a function of promoter architecture. Our observation that codon usage affects mRNA levels when we use fragments of the yeast genome shows that naturally existing variation in codon bias among native yeast genes directly affects mRNA expression. The fact that our predictions of mRNA levels are not perfect (R^2^ < 1) means that there is still much to be understood regarding the exact molecular mechanisms by which this occurs. The storage of public data and metadata in a standardized format allowed us to identify the mechanism by which the effect of sequences in the open reading frame are modulated by promoter architecture. Overall, the better and more quantitative data become, the more clear it is how little in gene regulation we truly understand (Cohen 2017).

Genetic screens involving targeted and random mutation in both *cis* and *trans* regulatory elements have played a major role in defining the machinery that controls gene expression (Struhl 1995; Sharon et al. 2012). However, the global output of all biomedical research is exponentially greater than the ability of any single lab. In the genomic era, naturally existing genetic variation has been utilized to understand gene regulation (Rockman and Kruglyak 2006; Lindeboom et al. 2016). In this era of publically available data we are rapidly approaching a point at which functional genomic data will be available for almost all mutants in a wide variety of conditions. Here we show that publically available expression data can be used to discover molecular mechanisms underlying entirely novel types of gene regulation.

How gene expression is regulated was recognized as a problem prior to the identification of DNA as the genetic material (Monod 1966) and reviewed in (Monod 1947). Understanding how expression levels are encoded in the genome and how genetic variation affects expression has been one of the biggest challenges in molecular biology for the past fifty years (Beer and Tavazoie 2004; Rockman and Kruglyak 2006). Much of this work has focused on non-coding regions, specifically on the ability of transcription factors to bind distinct regulatory motifs in promoters and enhancers. The limited work that has been done understanding the regulatory potential of coding sequences has been done in an isolated manner, without considering the other regulatory regions of the gene (Kudla et al. 2009; Puchta et al. 2016; Radhakrishnan et al. 2016). Our results here show that coding sequences contain a large amount of regulatory information that affects gene expression, and that this information interacts with information in the promoter architecture. Experimental systems and computational analyses that treat regulatory units in isolation miss molecular mechanisms that affect the expression of thousands of genes.

Why do organisms implement regulation at the level of the ORF? Measurements of steady-state mRNA levels from cells proliferating at different rates suggested that thousands of genes are regulated in response to changes in growth rate in both yeast (Brauer et al. 2008) and mammalian cells (Badia et al. in progress). However, promoter activity, as measured by promoter-YFP signal, scales with growth rate similarly for most genes, suggesting little or no promoter-specific regulation with growth rate (Keren et al. 2013). Here we show that information encoded in the ORF and interpreted in a manner that depends on the promoter architecture and on the RNA helicase Dbp2, resolves this conflict. Constitutive genes, which are the majority of genes in yeast, are regulated mostly by sequences encoded in the ORF. Why? Translation-mediated regulation is less noisy than that mediated by transcription factors (Carey et al. 2013). Furthermore, it may be mechanistically easier to make coordinated quantitative changes in the mRNA levels of thousands of genes in a post-transcriptional manner, compared to through the use of transcription factors. Therefore implementing global regulation at the level of the ORF may be a more reliable and efficient means to coordinate changes in expression across thousands of genes.

To implement biologically useful regulation via coding sequences, the regulation needs to be targeted to some genes and not others. One way is to have large sequence composition differences between groups of genes. A second way is to somehow insulate some genes but not others from this mechanism of regulation. Dbp2 is well positioned to do the latter. As an RNA helicase it is likely to be relatively sequence-neutral compared to many RNA binding proteins. It remains to be identified if Dbp2’s helicase property enables regulation based on differences in secondary structure, as was recently observed for Dhh1 (Jungfleisch et al. 2017). In addition, Dbp2 associates with chromatin in a nascent-RNA dependent manner and is co-transcriptionally loaded onto mRNAs (Cloutier et al. 2012; Ma et al. 2016). Dbp2 is required for multiple stages of co-transcriptional mRNP assembly. This raises the possibility that Dbp2 specificity is encoded in the promoter via selective loading of Dbp2 onto certain classes of transcripts, such as those with TATA boxes. This would in turn regulate mRNP assembly and nuclear export. Dbp2 co-IPs with RPL2A (ribosomal 60S subunit protein L2A), TIF4631 (the translation factor eIF4G) the SEC 13 (nuclear pore complex) (Gavin et al. 2002), suggesting that Dbp2 may remain bound to the mRNA throughout its lifecycle, directly linking transcription initiation with translation.

## Methods

### Library plasmids construction

All plasmids are named and described in **Supplemental Table 3**. To generate the random gDNA fragment library we used as a vector backbone the plasmid pRS416 carrying the URA3 selectable marker. We cloned into pRS416 the GALL inducible promoter and CYC1 transcriptional terminator, generating P47. The GALL promoter is a version of the GAL1 promoter truncated to remove the two most distal Gal4 binding sites, and has approximately 10% of the promoter activity as the fully intact GAL1 promoter(Mumberg et al. 1994). We next used Gibson assembly (NEB, E2611S) to insert the ATG-ClaI-Stop fragment (**Supplemental Fig. 1A**). This fragment was comprised of two annealed oligonucleotides, 174 and 175 (see **Supplemental Table 4**) in 10X Oligo Annealing Buffer (400 μl 1M Tris pH 8, 80ul 0.5M EDTA pH8, 800ul 2.5M NaCl, 2720 μl DNase/RNase-Free Water) in a reaction containing: top and bottom strand DNA oligos (200 μM each), 2 μl of 10X Oligo Annealing Buffer and DNase/RNase-Free Water for a total volume of 20 μl After incubation at 94°C for 4 minutes, we allowed the reaction to cool to room temperature for 10 minutes. To obtain the RPL4Apr plasmid (P86) we amplified using Phusion polymerase (Thermo Fisher Scientific, F530S) in GC buffer the RPL4A promoter from genomic DNA using primers 222 and 223 (**Supplemental Table 4**) and cloned the amplicon via Gibson assembly (NEB, E2611S) into the SacI and XbaI restriction sites of plasmid P47.

### Library construction

The random yeast genomic sequences were obtained by cutting 40 μg of isolated genomic DNA (MasterPure^TM^ Yeast DNA Purification Kit. Epicentre, MPY80200) from yeast strain FY4, a prototrophic S288C strain, for 4 hours at 37 °C with a combination of 4 restriction enzymes (TaqαI, HpaII, MspI and AciI). All restriction enzymes used in this study are from New Englands Biolabs. Size selection was performed using a 2.5% agarose gel and purified DNA was ligated into ClaI cut and dephosphorylated (rSAP, New Englands Biolabs, M0371S) vector (P47). Ligated products were desalted by drop dialysis using 13 mm diameter, Type-VS Millipore membrane (Merck Millipore, VSWP01300) and 3 μl were transformed by electroporation using the UltraClone Kit with 10G Elite DUO (Lucigen, 60117-1). *E. coli* transformants were selected on LB medium (0.5% yeast extract, 1% NaCl, 15% bactotryptone) supplemented with 100 μg/ml ampicillin as a selection marker.

To obtain the RPL4Apr plasmid library with the same inserts that are present in the GALLpr library we amplified with Q5 DNA polymerase reaction mix (NEB, M0491S) the library previously cloned into P47 using primers 345 and 198 (see **Supplemental Table 4**). After cutting pRS416- RPL4Apr (P86 in **Supplemental Table 3**) with XbaI and XhoI restriction enzymes we cloned the PCR library fragments into pRS416-RPL4Apr via Gibson assembly (NEB, E2611S), obtaining the new promoter library that was transformed in *E. coli* electrocompetent cells (UltraClone Kit with 10G Elite DUO. Lucigen, 60117-1) and plated on LB medium (0.5% yeast extract, 1% NaCl, 15% bactotryptone) supplemented with 100 μg/ml ampicillin as a selection marker. The constitutive and inducible library plasmids were purified using the Nucleobond XtraMidi Plus EF (Machery-Nagel, 740422.50).

### Yeast transformation

The gDNA libraries cloned into P47 (GALL) and P86 (RPL4A) were transformed into BY4741 (Y49) ( **Supplemental Table 3**) via the lithium acetate method(Gietz and Woods 2006) and plated in glucose synthetic complete dropout plates lacking uracil for plasmid selection. In the same way, we transformed the GALL inducible library of the yeast native ORF in the mutant strains Y194 and Y195 ( **Supplemental Table 3**). After 2 days of growth at 30C we collected all the transformants and we resuspended them to OD of 0.5 in synthetic complete dropout plates lacking uracil supplemented with 2% galactose as a carbon source in order to get the GALL promoter induction. After growing the cells overnight we scraped off the plates and inoculated four 500 ml flasks of the same medium to OD of 0.025 as biological replicates. The cultures were grown about 10 hours at 30°C to reach an exponential phase with an OD of 0.5, where we took 1.5 ml samples in order to extract plasmid DNA (MasterPure^TM^ Yeast DNA Purification Kit, Epicentre) and transcribed RNA (MasterPure^TM^ Yeast RNA Purification Kit, Epicentre) from all the biological replicas. We pelleted at 4 °C and froze every sample at −80°C until processing.

### NGS sample preparation

We used a gene specific primer to generate the cDNA from each RNA preparation (**Supplemental Fig. 1**, primer 198 depicted in yellow and see **Supplemental Table 4**) using the ThermoScript RT-PCR System (Life Technologies, 11146-024). With barcoded primers we amplified the second DNA strand and the library plasmids DNA in order to differentiate between replicas and sample origin, in this case, DNA and RNA (for details of barcoded primers combination for the libraries see **Supplemental Table 4 and Supplemental Table 5**). We note that because we made cDNA using a primer in the CYC1 terminator, any insert that results in transcription termination would give little or no mRNA expression in our assay. The PCR was set up as 50 μl reaction of Q5 DNA polymerase (NEB, M0491S) in the following cycling conditions: 98°C for 30s; 98°C for 10s, 55-64°C (depending on primer combination for demultiplexing on technical variability and biological replicates) for 30s and 72°C for 30s (20 cycles); and 72°C for 2 min. PCR products were purified using MinElute PCR Purification Kit (Qiagen, 28004). NGS libraries were prepared from 100 ng of the purified DNA amplicons using Ovation Rapid DR System (Nugen, 0319-32) according to manufacturer's instructions. Each library was visualized on a Bioanalyzer (Agilent Technologies) and quantified by qPCR with a Kapa Library Quantification Kit (Kapa Biosystems, KK4835). The libraries were sequenced through Illumina HiSeq 2500 125bp paired-end reads method.

### NMD reporter plasmids construction and measurement

The four NMD reporter constructs GALLpr-mCherry-PTC-YFP, GALLpr-mCherry-linker-YFP, RPL4Apr-mCherry-PTC-YFP and RPL4Apr-mCherry-linker-YFP (P71, 68, 74 and P78, respectively. For details see **Supplemental Table 3**) were generated by fusion PCR reaction with overlapping oligos amplifying mCherry with primers 538 and 539 (in case of stop codon presence in the linker between mCherry and YFP) or primers 538 and 540 (in case of stop codon absence in the linker between mCherry and YFP). YFP was amplified with primers 541 and 542 (see **Supplemental Table 4** for primers details). For all PCR reactions Phusion polymerase (Thermo Fisher Scientific, F530S) in GC buffer was used. After fusion of both fragments, the final product was purified using MinElute PCR Purification Kit (Qiagen, 28004) and cloned in the ClaI cut and dephosphorylated with rSAP (New Englands Biolabs, M0371S) pRS416-GaLLpr and pRS416-RPL4Apr vectors (P47 and P86 in **Supplemental Table 3**). The four NMD reporter plasmids were transformed into Y49 and multiple transformants of each construct were inoculated into 96 well plates containing SCGal-URA media. Cells were grown overnight, diluted 1:50, and measured by flow cytometry after 15 hours of growth, when the OD600 was around 0.5. Wells in which the cell density was too high or too low were discarded. All flow cytometry was performed on a BD LSRFortessa (BD Biosciences) with 488nm and 561nm lasers with 530/28 and 610/20 filters for YFP or mCherry, respectively. Analysis of flow-cytometry data was performed using a previously described custom MATLAB pipeline (Carey et al. 2013). These two promoters have approximately equal expression levels (**Supplemental Fig. 4**). The same procedure was done for the constructs in Y194 and Y401 strains.

### Sequencing data processing

Samples and replicates for the library were all compact in fastq files with 125bp paired-end reads. The first step in processing the reads was removing 3’ Illumina adapters of those reads where the inserted ORF was smaller than 125 bp. Next, we took advantage of the oligos used during the PCR for the sequencing library preparation to demultiplex each sample and replicate (split reads by type of sample origin - DNA or RNA - and replicates 1 or 2). Then the oligos were trimmed from the read using cutadapt (Martin 2011), leaving part of the unique genomic DNA sequence that was inserted in the plasmid as an ORF. To obtain the entire gDNA sequence inserted, we mapped all trimmed reads to the yeast genome using bowtie2(Langmead and Salzberg 2012). Hits where the forward and reverse reads mapped not uniquely, in different chromosomes, same strand, or the distance between them was >1Kb, were directly discarded. The resting hits were used to obtain the complete gDNA fragment used in the library and the number of reads mapped used as a quantification measure of number of DNA or RNA molecules. For downstream analysis we attached the 5’ and 3’ constant flanking sequence of the plasmid (which contained the designed ATG and stop codons) for later quantify other features such as 3’UTR length (measured as the length from the first stop codon in the ORF to the start of the CYC1 transcription terminator**, Supplemental Table 6**).

### Expression values and data analysis

After obtaining the number of reads per unique sequence for each sample and replica we first normalize read counts by the total amount of reads per sample and replica using the formula:

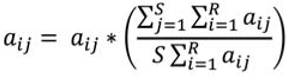

Where a_ij_ represents the read count for a unique sequence variant per replica and sample. S is the number of samples (RNA, DNA and technical replicas) and R the total number of unique transcripts. In order to normalize by technical and biological bias we used as values of expression the log_2_ ratio between the averaged replicas DNA

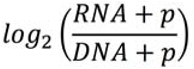

reads and RNA reads:

Where p represents a pseudocount (p = 0.5). In order to uniformize the measurements across all libraries we calculated expression as the z-score value of this last log_2_ ratio.

It is possible that the effect of codon bias may be partially co-transcriptional; there is good experimental evidence that GC content affects the rate of mRNA synthesis, and limited evidence that codon bias itself does so(Newman et al. 2016; Zhou et al. 2016). To avoid possibly confounding effects we regress out the contribution of insert length and GC content (see methods). This allowed us to measure the GC and length-independent role of 3’UTR and tAI to expression.

### Model to predict expression in the ORF library using inserts features

To build a multiple linear regression model to predict expression values based on the composition of the ORF (**Supplemental Fig. 3** illustrates all steps followed) we used already described transcript features (tAI, 3’UTR length, GC content and transcript length) and others of unknown mechanism. From the subset of fragments without PTCs that are fully contained within native yeast ORFs, the directionality of the inserted fragment had a significant impact on expression (fragments oriented in the same direction as the native gene had higher expression that reverse oriented ones, t-test p<10^−11^). Since the genomic orientation of the inserted ORF affects expression but the mechanism is unknown, we used a machine learning approach to identify the ORF features that better discriminate between forward or reverse oriented sequences to later include them in the final model to predict expression. We built three different classifier models (Logistic regression, Naive Bayes and Tree Bagger) based on > 4000 sequence features (bases counts, amino acid counts, codon counts, frequency of nts in different position of codons, hexamers…). All three classifiers were trained on 1/3 of the data and the number of features of each model was reduced to dozens by sequential feature reduction (‘sequentialfs’ Matlab function). The three classifiers performed correctly (median AUC = 0.80) when tested in the remaining 2/3 of the dataset. To determine what features could predict expression we took all the selected features from the classifiers and used them in a linear model to predict expression. Using LASSO, to further reduce the number of predictors, we obtained 5 sequence features that could explain 11% of the variance in expression: frequency of A or G in the first position of the codon (AG1), A or C in the second position of the codon (AC2), frequency of A in the second position of the codon (A2) and the frequency of appearance of two the hexamers GAAAGA and ACGTTA. In the final model to predict expression of all variants in our library we included the three features AG1, AC2 and A2, as these are likely to be functionally interpreted by the translating ribosome inside the cell. For all model predictions we used a 10-fold cross-validation scheme to assess the accuracy of all models and to test for overfitting.

### Predicting mRNA level of native genes based on ORF features

To measure the contribution of the coding region to changes in expression we used a linear regression model that included exclusively ORF features selected previously using the gDNA library. 3’UTR length and premature termination codon features were not used for models for native genes. We used 10-fold cross validation to test the accuracy of the model. mRNA expression levels are from (van Dijk et al. 2015). All mRNA-seq datasets give essentially the same result; this can be seen in (**Fig. 3B**), which shows results from a more limited ORF sequence feature model that includes just a single predictor. The same model was used to predict the expression of genes for which the promoter contribution to expression was previously measured (Keren et al. 2013).

### Yeast NMD strength analysis

To measure NMD strength for native yeast genes wild-type (SRR1258470,SRR1258471) and *upf1* (SRR1258533,SRR1258534) RNA-seq reads were downloaded from NCBI SRA (PRJNA245106) (Smith et al. 2014). Expression was quantified using kallisto (Bray et al. 2016) (--single -l 180 -s 20) and orf_coding_all.fasta from SGD (Cherry et al. 2012) (yeast genome R64-2-1). For each gene the NMD effect was defined as log2( mean(expression *upf1*) / mean(expression WT)). To work in a parameter regime comparable to that of the gDNA library, only genes with 3’UTR lengths (Nagalakshmi et al. 2008) from 200 - 400nt were analyzed.

### Digital expression explorer analysis

We obtained the processed data of 6402 RNAseq dataset from the digital expression explorer database (Ziemann et al. 2015). Expression was calculated normalizing sequence read counts by transcript length and sum of reads in each experiment. The effect of codon bias on expression was inferred for each experiment as described in Results (**Fig. 2F**). To ensure a meaningful calculation of the effect of codon bias (that assumes Promoter-YFP data (Keren et al. 2013) to be correlated with mRNA levels) we removed experiments with a low correlation between expression and Promoter-YFP data (correlation < 0.4). We next searched for experiments in which the promoter-dependent effect changes from the “wild-type” behavior. We grouped the expression datasets by experiment (BIOPROJECT) (each setup corresponds to a different work) and graphed each of them separately in the same space (TATA/TATA-less vs “effect of codon bias on expression”). We manually looked for BIOPROJECTS in which a subset of similar samples (e.g., biological replicates of a mutant strain) differs consistently from another subset (e.g., wild-type strains).

### Data access

DNA-seq and RNA-seq data generated in this study are deposited in the National Center for Biotechnology Information Gene Expression Omnibus under accession GSE100452. All code for figures and analysis are on GitHub (https://github.com/MikiSchikora/PromoterarchitectureEspinar17/).

## Acknowledgments

L.B.C. was supported by Ministerio de Economía y Competitividad (MINECO) (BFU2015-68351-P) and AGAUR (2014SGR0974). We’d like to thank Wenfeng Qian, Shaohuan Wu, Aaron New, Francesc Posas, and members of the Barcelona Yeast Group and the CRG/UPF Computational Genomics and Systems Biology groups for helpful comments on the results and manuscript.

## Supplementary Figures

**Supplemental Fig. 1.**
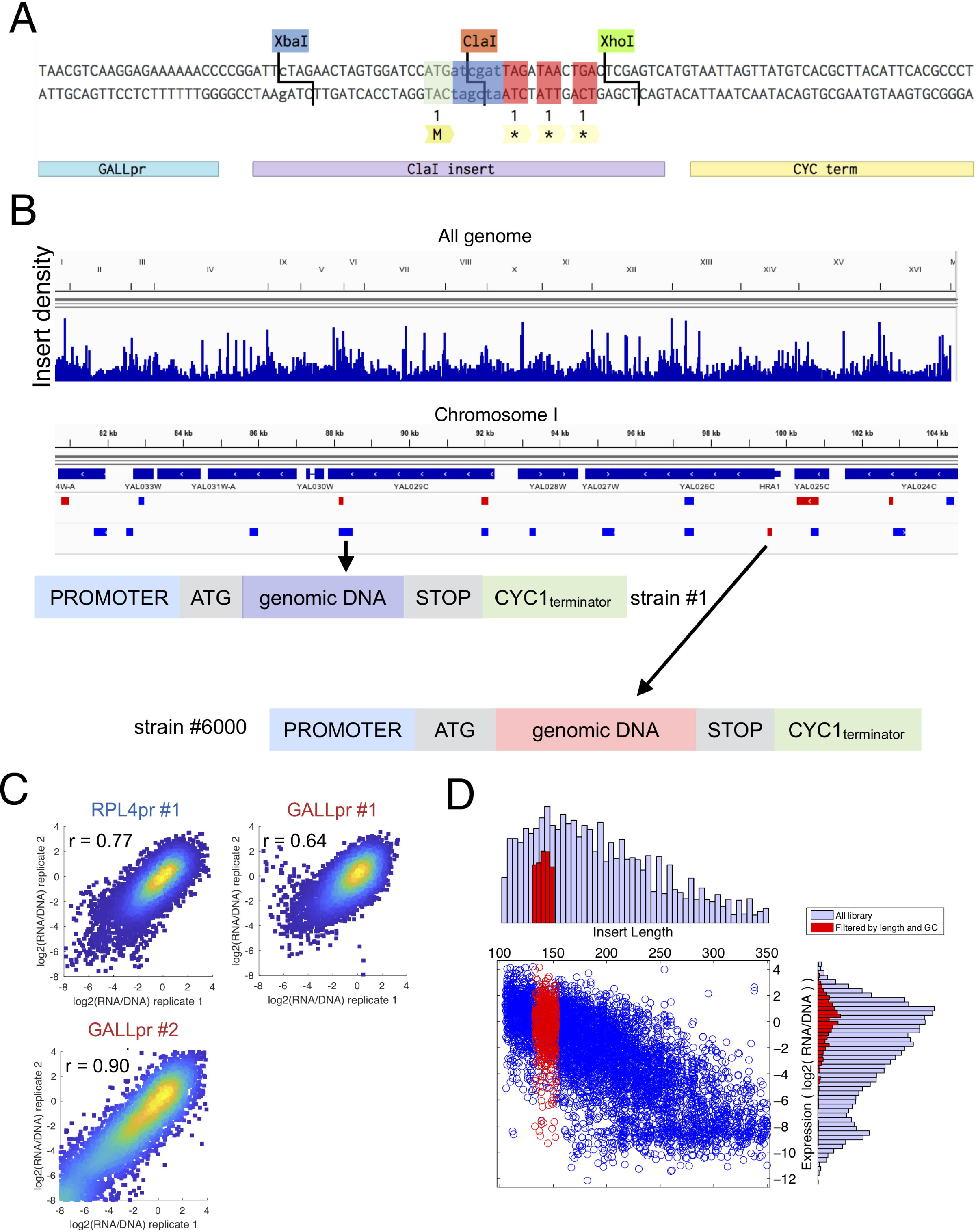
gDNA library and construct. (**A**) Sequence details of the ClaI restriction site (dark blue) where fragments of the yeast genome are cloned. The construct contains a GALL promoter (left-bottom blue) a designed region where the ClaI site is located (bottom purple annotation) and the CYC1 terminator (bottom yellow annotation). The designed region contains a start codon ATG (green), followed by the ClaI site and finishing with three stop codons (red) separated by one nucleotide to cover all possible reading frames. (**B**) Top: distribution of the genomic fragments inserted in the GALL library. Bottom: two example resulting strains after insertion of two chromosome I fragments. (**C**) Expression measurements across replicates are reproducible. Shown are two independent creations of the GALL promoter library, and one of the RPL4A library. We note that if differences in signal (experimental error) were responsible for the results, then we would expect the codon bias and NMD effects to be stronger in the experiments with a higher correlation between replicates. The opposite is true; both signals are also higher in the RPL4A library than in the GALL library #2 (see **Supplemental Fig. 5**). (**D**) Shown are the length and expression distribution for the entire GALL library (blue) and a small subset with similar insert length (133-153 nt) and GC content (0.35-0.45% GC). In this subset with expression varies by 4 orders of magnitude.

**Supplemental Fig. 2.**
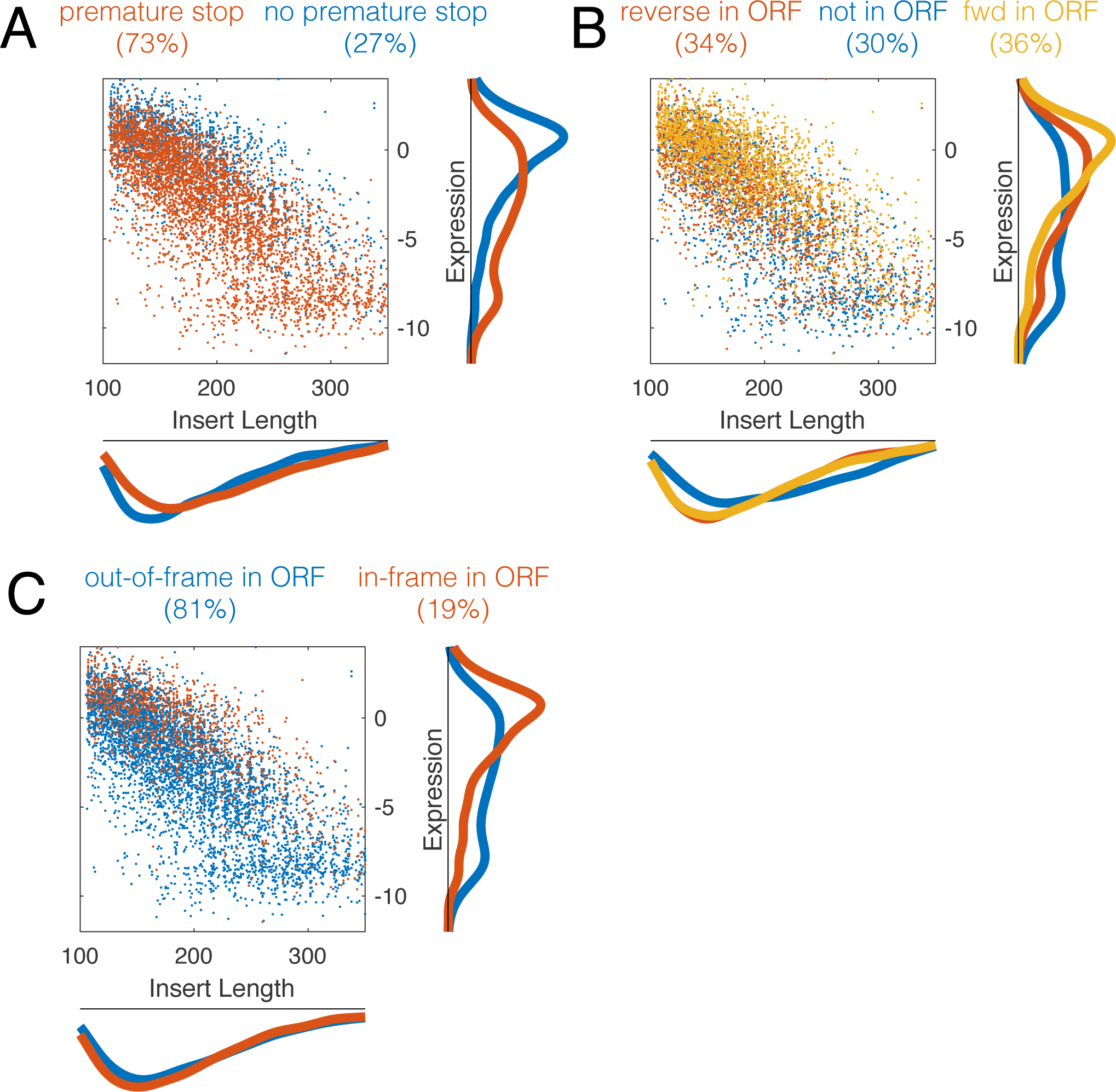
Breakdown of expression by insert origin. Inserts were classified according to (A) absence/presence of a PTC, (B) if they came from within an ORF or not, and (C) if the translational reading frame in our system is the same as the reading frame in the native gene.

**Supplemental Fig. 3.**
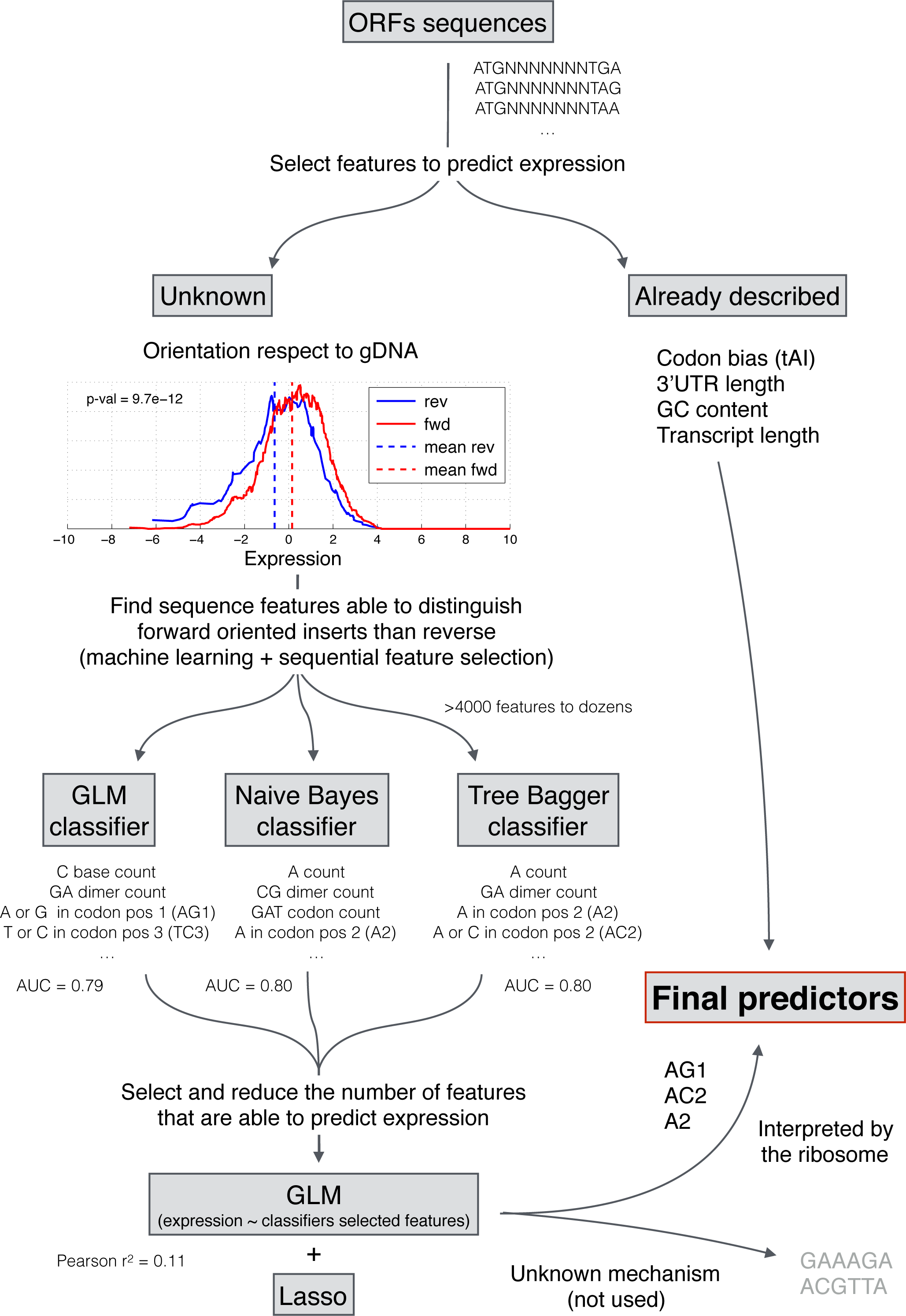
Flowchart of the steps followed to select the predictors of the ORF- expression model. AUC = Area Under the Curve, GLM = Generalized Linear Model, Lasso = Least absolute shrinkage and selection operator, AG1 = Containing an A or a G in codon position 1, TC3 = Containing a T or A in codon position 3, A2 = Containing an A in codon position 2, AC2 = Containing an A or C in codon position 2.

**Supplemental Fig. 4.**
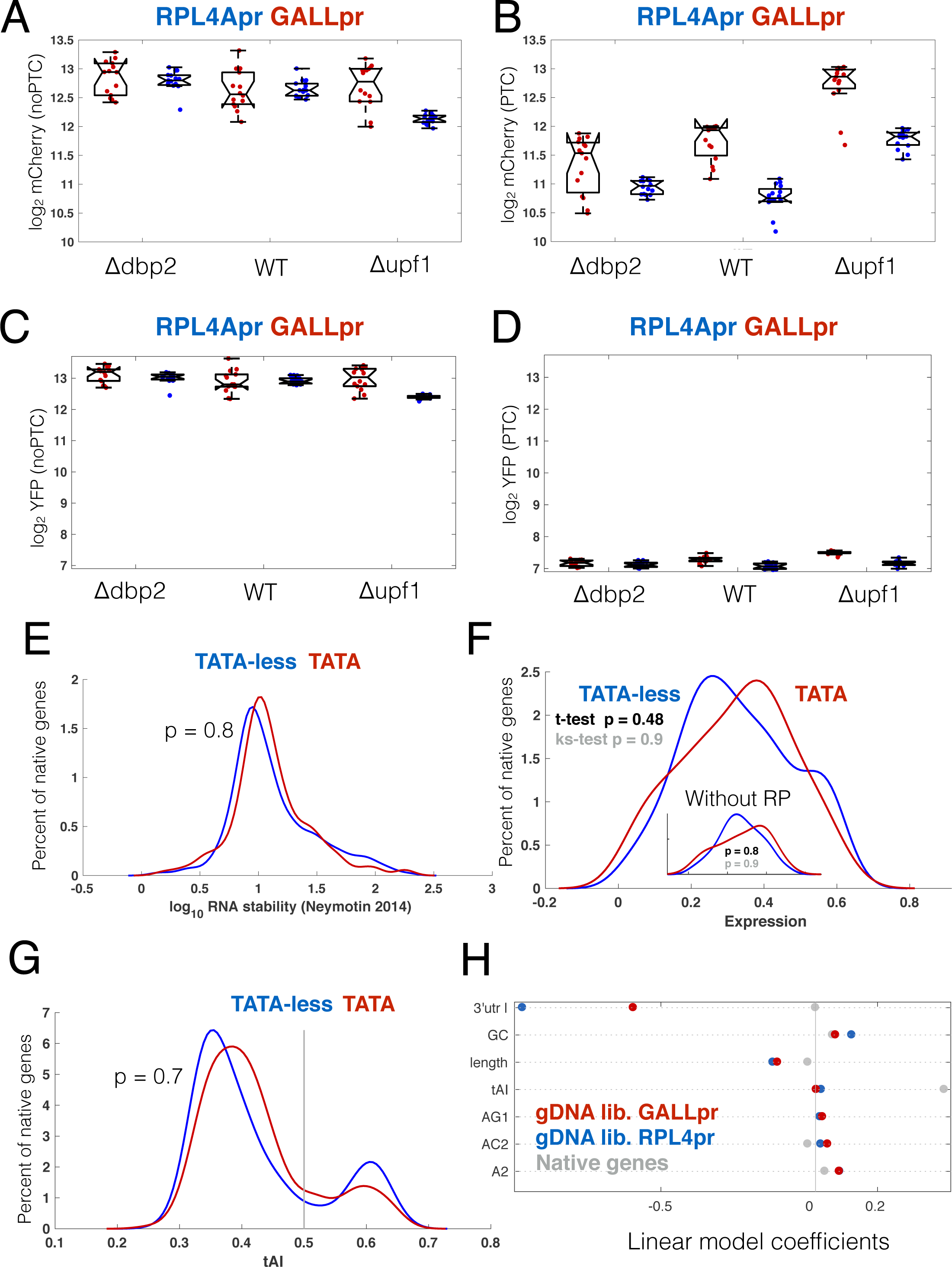
Several parameters of constitutive and regulated genes. (**A**) mCherry expression of the NMD reporter constructs lacking a premature termination codon (PTC) between mCherry and YFP (see **Fig. 3D**), driven by the constitutive (blue) and regulated (red) promoter. Each construct was measured in wild-type, *upf1Δ* and *dbp2Δ* strains. Each point is from a different biological replicate. (**B**) The same as in (A), but for constructs bearing a PTC in the intervening mCherry-YFP linker. (**C,D**) The same as (A,B), but showing YFP expression measurements. (**E**) Distribution of RNA half-life values reported Neymotin et al (Neymotin et al. 2014). Shown are these values for TATA (red) and TATA-less (blue) genes. The p value corresponds to a ttest. (**F**) Distribution of mRNA expression levels (in log_2_ TPM) for the same groups as in (E). The inset shows the expression distributions after removing ribosomal proteins. (**G**) tRNA Adaptation Index (tAI) scores for for the same groups as in (E). The line refers to the threshold used in the analysis of how tAI affects expression in native genes (see Results). (**H**) Shown are all of the coefficients for features used in the linear model that predicts expression from sequence features. 3’UTR length for native genes is zero as only features in the ORF are considered.

**Supplemental Fig. 5.**
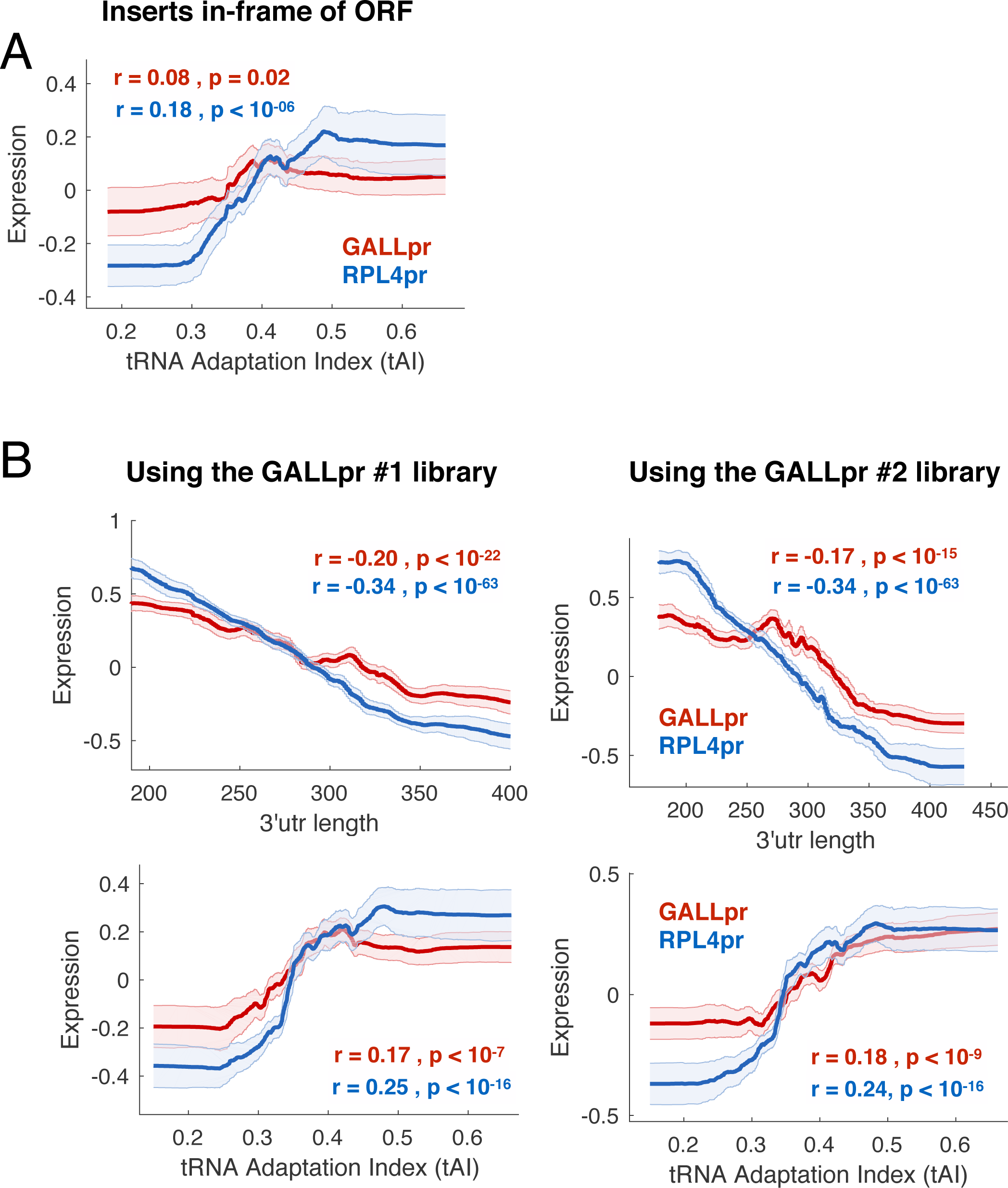
The tAI effect of in-frame fully-contained fragments and reproducibility across replicates of the gDNA library. (**A**) Shown are the correlation of tAI on insert length and GC content corrected expression for only the subset of inserts that lack a premature stop codon and that will be translated in the same reading frame in our library and in the native gene. (**B**) The RPL4A promoter library compared to the two GALL promoter libraries. While the two GALL promoter libraries have different correlations among replicates, the result is the same in both libraries.

**Supplemental Fig. 6.**
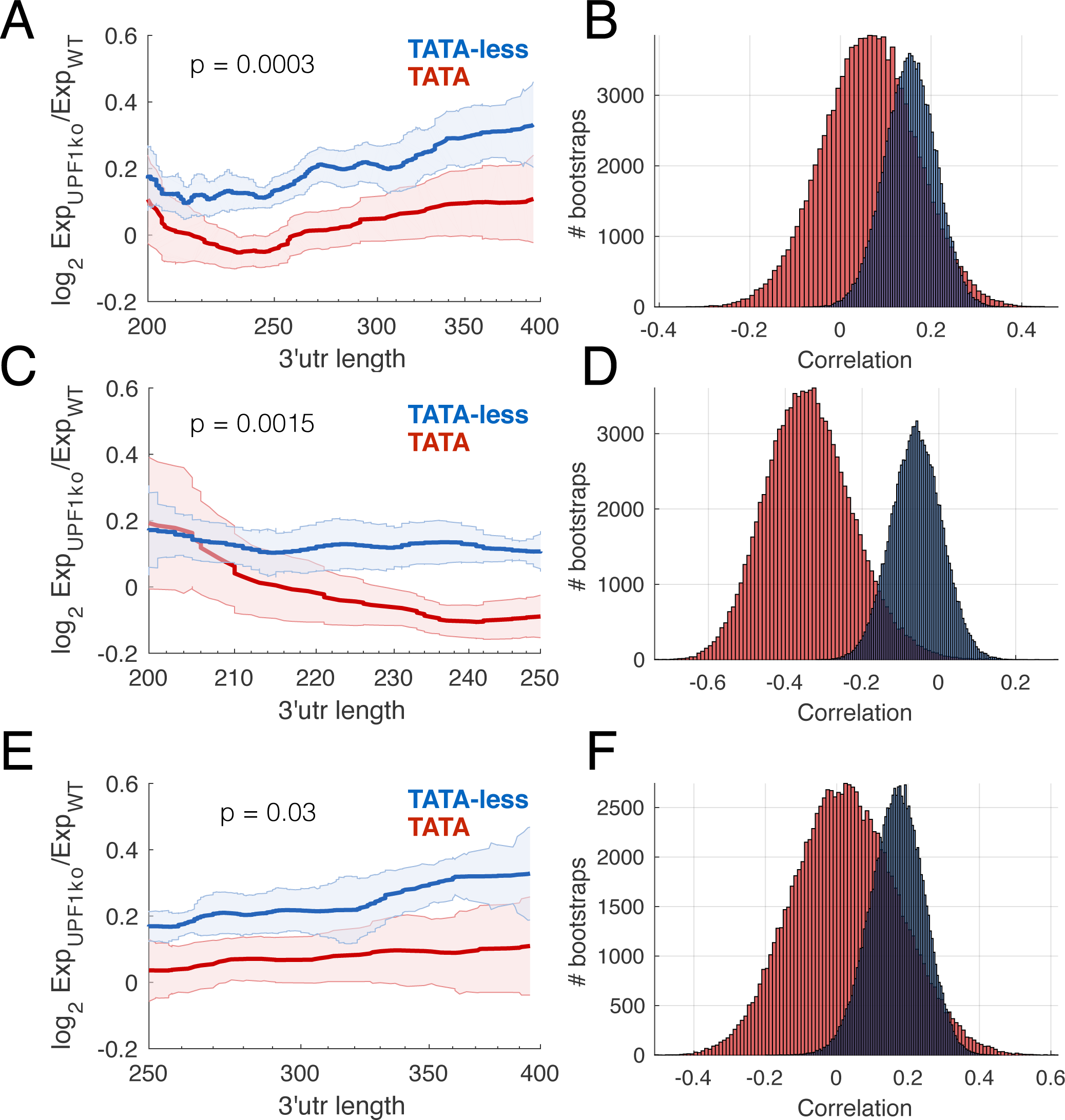
TATA-less genes are more affected by NMD at native transcripts with both short 3’UTRs and long 3’UTRs. (**A,B**) NMD strength appears to have two regimes, transcripts with a 3’UTR length less than 250nt, and those with a longer 3’UTR. (**C,E**) TATA-less transcripts have a stronger NMD effect (p-value for a t-test of mean NMD effect) for transcripts with both short and long 3’UTRs. (**B,D,F**) Correlation between 3’UTR length and NMD strength for calculated from bootstrapped sampling of transcripts.

**Supplemental Fig. 7.**
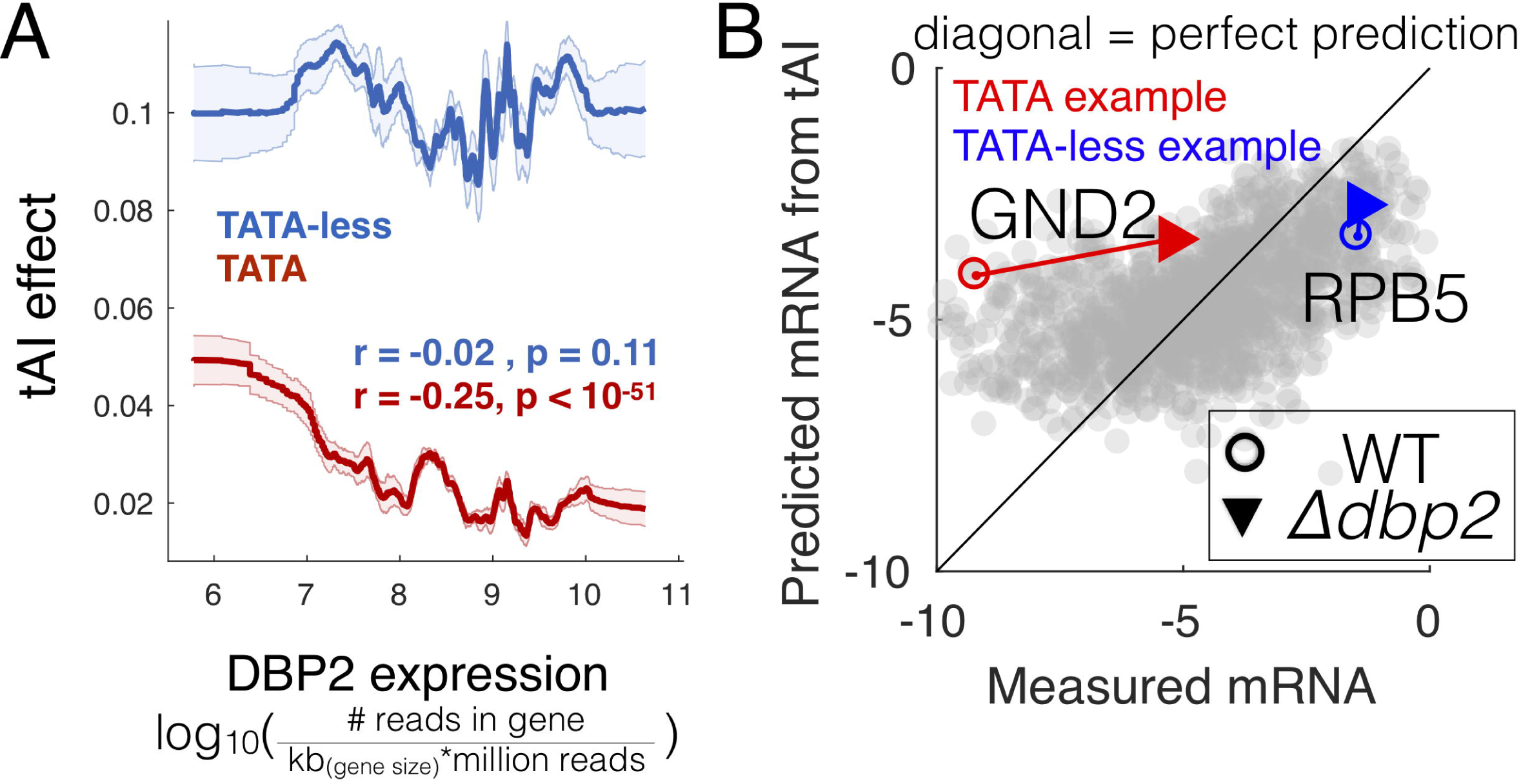
Low DBP2 expression correlates with a high tAI effect in TATA+ but not TATA-less genes across all DEE datasets. (**A**) The calculated the tAI effect for all RNA-seq datasets in yeast compared with the measured DBP2 expression. Lines show the median tAI effect and shaded error bars the standard deviation from bootstrapping. Experiments are binned by DBP2 expression. Consistent with all our previous data, conditions and mutants with low DBP2 expression have a higher tAI effect for TATA+ genes and no change for TATA-less genes. (**B**) Shown is the ability of tAI to predict steady-state mRNA levels for an RNA-seq experiment performed in a WT genotype (Beck et al. 2014). Two example genes with different promoters (TATA, and TATA-less) show different changes regarding the effect of tAI in a *dbp2Δ* strain, performed under the same conditions.

**Supplemental Fig. 8.**
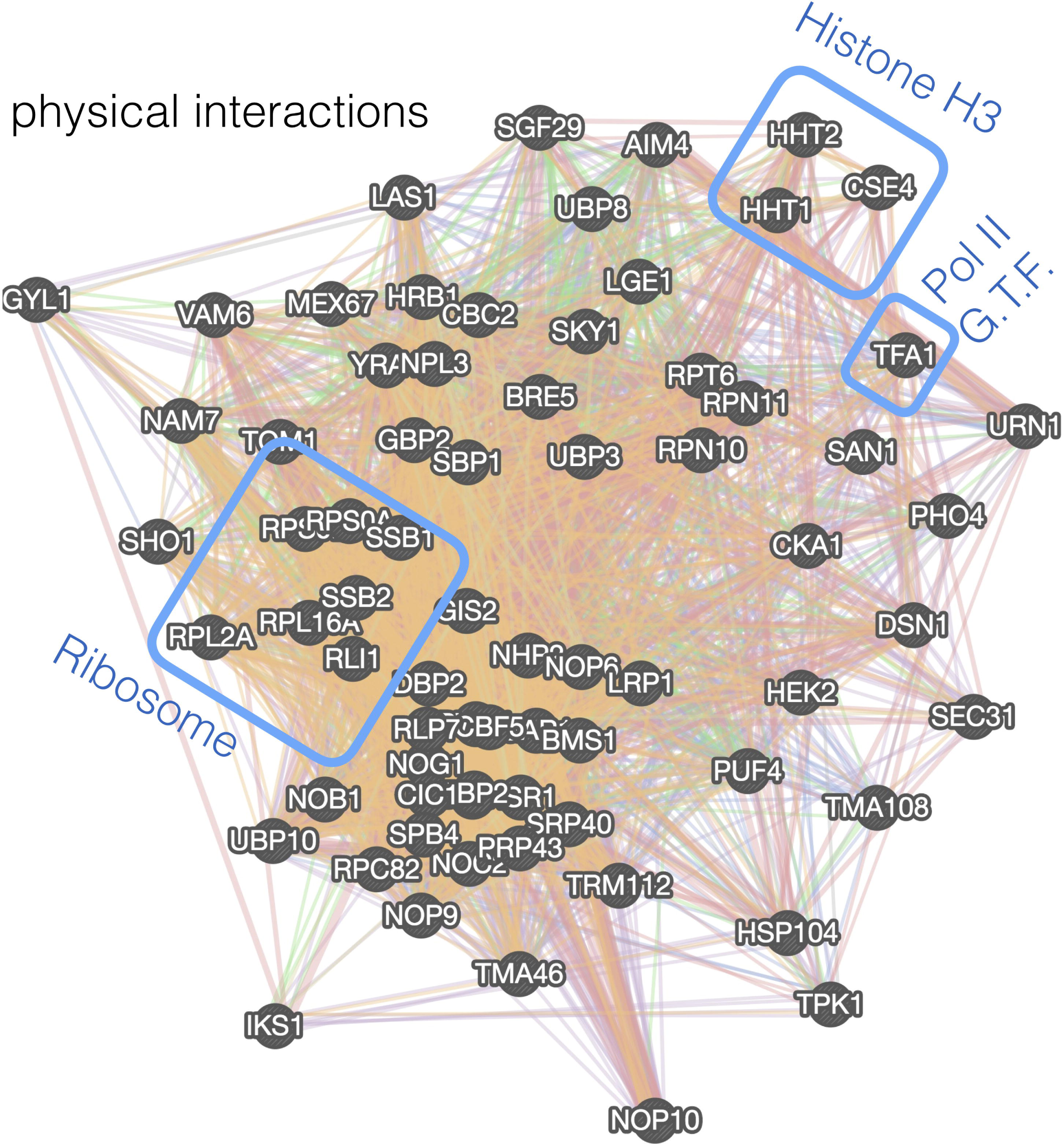
Dbp2 physically interacts with general transcription factors, chromatin, nuclear export machinery and translational machinery. (**A**) Dbp2 physical interactome of proteinprotein interactions (Cherry et al. 2012). Dbp2 interacts with ribosomal, histone H3 and General Transcription Factors (GTFs) of Pol-II (TAF1 is the yeast ortholog of the mammalian TFIIE).

### Supplementary Tables

**Supplemental Table 1**: Properties of the inserts in the ORF library

**Supplemental Table 2**: Results of Gene Ontology Analysis

**Supplemental Table 3**: Table of strains and plasmids used in this study

**Supplemental Table 4**: Table of used primers

**Supplemental Table 5:** PCR Primer barcode combination

**Supplemental Table 6**: Nucleotide sequences of 5′ and 3′ regions of random ORF library plasmid construct (5′ to 3′)

